# Impact of homologous recombination on core genome evolution and host adaptation of *Pectobacterium parmentieri*

**DOI:** 10.1101/2023.12.15.571927

**Authors:** Dario Arizala, Mohammad Arif

## Abstract

Homologous recombination is a major force mechanism driving bacterial evolution, host adaptability and acquisition of novel virulence traits. *Pectobacterium parmentieri* is a plant bacterial pathogen distributed worldwide, primarily affecting potatoes, by causing soft rot and blackleg diseases. The goal of this investigation was to understand the impact of homologous recombination on the genomic evolution of *P. parmentieri*. Analysis of *P. parmentieri* genomes using Roary revealed a dynamic pan-genome with 3,742 core genes and over 55% accessory genome variability. Bayesian population structure analysis identified seven lineages, indicating species heterogeneity. ClonalFrameML analysis displayed 5,125 recombination events, with the lineage 4 exhibiting the highest events. FastGEAR analysis identified 486 ancestral and 941 recent recombination events ranging 43 bp - 119 kb and 36 bp - 13.96 kb, respectively, suggesting ongoing adaptation. Notably, 11% (412 genes) of the core genome underwent recent recombination, with lineage 1 as the main donor. The prevalence of recent recombination (double compared to ancient) events implies continuous adaptation, possibly driven by global potato trade. Recombination events were found in vital cellular processes (DNA replication, DNA repair, RNA processing, homeostasis, and metabolism), pathogenicity determinants (type secretion systems, cell-wall degrading enzymes, iron scavengers, lipopolysaccharides, flagellum, etc.), antimicrobial compounds (phenazine and colicin) and even CRISPR-Cas genes. Overall, these results emphasize the role of homologous recombination in *P. parmentieri*’s evolutionary dynamics, influencing host colonization, pathogenicity, adaptive immunity, and ecological fitness.

**Significance:** This study explores the influence of homologous recombination on the genomic evolution of the globally distributed plant pathogen *P. parmentieri*, characterized by highly heterogenous strains from various global locations. Our findings reveal diverse recombinogenic patterns within the core genomes of *P. parmentieri* isolates, notably in genomic loci associated with vital cell functions, pathogenicity determinants, and CRISPR-Cas genes. These findings highlight the role of homologous recombination in shaping the genomes of *P. parmentieri* and impacting its phytopathogenic lifestyle. Additionally, the data suggest a potential role of recombination in the ecological adaptation of this species across different climates, providing insights into the worldwide presence of *P. parmentieri*. This study represents a pioneering exploration of the impact of homologous recombination on the dynamic evolutionary genomics of the soft rot-causing bacterium *P. parmentieri*.

## Introduction

In various organisms, including plant bacteria, homologous recombination constitutes an essential genetic mechanism (González-Torres et al. 2019). Plant bacteria engage in homologous recombination through a process known as transformation, enabling them to uptake foreign DNA from the surrounding environment. The process involves the exchange and recombination of genetic material between two similar DNA sequences, leading to the emergence of new gene combinations and potential changes in phenotype (González-Torres et al. 2019; Kuzminov 2011). For homologous recombination to occur, both ends of the recombined fragment must exhibit a high degree of DNA sequence similarity. As a result, this molecular mechanism primarily occurs between strains of the same species and taxonomically related bacteria (David et al. 2017; Jibrin et al. 2018; González-Torres et al. 2019). The need for a certain level of identity explains why the highest rates of homologous recombination are found in sections of the core genome where sequence similarities are preserved (Dixit et al. 2015; Everitt et al. 2014; Jibrin et al. 2018).

Homologous recombination in bacterial pathogens is of great importance for genetic diversity and adaptation, as it enables the acquisition of new traits and the evolution of these microorganisms. Through homologous recombination, bacteria can acquire new genes and alleles that confer advantages, such as the rapid spread of traits enabling pathogen survival, overcome host defense mechanism, and establishing successful infections (David et al. 2017; Jibrin et al. 2018; González-Torres et al. 2019). During the recombination process, bacteria can exchange genes involved in virulence, antibiotic resistance, and environmental adaptation, leading to the emergence of new pathogenic variants (Prokchorchik et al. 2020). Importantly, homologous recombination has been shown to accelerate evolution by up to a hundred times when different strains are present in the same host (Didelot et al. 2016). This genetic diversity helps plant bacterial pathogens in colonizing a range of hosts, evading plant defenses, and thriving in various environments (Vanhove et al. 2019).

In addition, homologous recombination functions as the main method for DNA repair in plant bacteria (David et al. 2017). It aids in repairing DNA damage caused by environmental factors, such as breaks or mutations (González-Torres et al. 2019). Plant bacteria may repair their genomes and maintain genomic integrity by exchanging genetic information with a homologous undamaged sequence (Potnis et al. 2019). This ensures their existence and aids in their ability to adapt to shifting environmental factors (Jibrin et al. 2018). Besides its role in DNA repair, recombination has also been observed to help in the deletion of deleterious mutations and the introduction of beneficial mutations during DNA synthesis, acting as a positive side effect leading to an energy source (David et al. 2017). Lastly, some studies have found that bacteria involved engage in homologous recombination to delete mobile genetic elements from their genomes (David et al. 2017). In summary, homologous recombination in plant bacterial pathogens is a key process that drives genetic diversity, adaptation, DNA repair, and the acquisition of new traits. Therefore, understanding the mechanisms and dynamics of this process can have profound implications for developing effective strategies to manage and control bacterial pathogens, as well as for optimizing crop health and productivity in agricultural settings.

*Pectobacterium parmentieri* is classified as a pectinolytic rod-shaped, facultatively anaerobic bacterium belonging to the genus *Pectobacterium* within the Pectobacteriace family (Adeolu et al. 2016). This species was formerly known as *P. wasabiae*, the horse radish pathogen, and later assigned as new species (Khayi et al. 2016). Like other members within the *Pectobacterium* genus, *P. parmentieri* is a plant-pathogenic bacterium causing soft rot and black leg diseases (Nykyri et al. 2012; Charkowski 2018; Zoledowska, Motyka-Pomagruk, et al. 2018). The bacterium is widely distributed across the globe, having been reported in several countries, including Belgium, France, Finland, Germany, Norway, Switzerland, The Netherlands, Canada, South Africa, New Zealand, Malaysia, Poland, Spain, Turkey, China and different states within the United States (Moleleki et al. 2017; De Boer et al. 2012; Ngadze et al. 2012; Pitman et al. 2010; Pasanen et al. 2013; Zoledowska, Motyka, et al. 2018; Arizala et al. 2019; Cao et al. 2021; Suárez et al. 2017; Ge et al. 2021). The strains of this species have been isolated from various environments and sources, including plants, soil, water, and plant debris. In a recent study, the metabolic modeling of the Finland strain SCC3193 revealed that *P. parmentieri* can adapt and survive in either soil or rhizosphere (Zoledowska et al. 2019). Although most *P. parmentieri* strains deposited in the NCBI GenBank database have been isolated mainly from potato (accessed on 14 September 2022), there are different surveys reporting the isolation of this bacterium from other hosts such as cabbage, tomato, eggplant, onions, carrot, maize, sugar beet, calla lily, sweet potato and star of Bethlehem, impacting both yield and quality (Zoledowska et al. 2018b).

The ability of *P. parmentieri* to cause soft rot diseases and establish a successful infection depends on the production of plant cell wall-degrading enzymes, namely cellulases, pectinases, and proteases (Nykyri et al. 2012; Arizala & Arif 2019; Zoledowska et al. 2018a). These enzymes are delivered via the type I (TISS) or II (TIISS) secretion systems, allowing the pathogen to macerate plant tissues, and spread throughout the host tissues (Nykyri et al. 2012; Zoledowska et al. 2018a; Arizala & Arif 2019; Arif et al. 2022). Importantly, due to the absence of the type III (T3SS) secretion system, studies have reported that *P. parmentieri* might be less virulent compared to other *Pectobacterium* species (Kim et al. 2009; Nykyri et al. 2012; Arizala & Arif 2019). However, other research studies have found highly pathogenic *P. parmentieri* strains on potato plants and tubers (Ge et al. 2021), demonstrating that the lack of T3SS in this bacterium is not a detrimental pathogenicity determinant for disease development (Kim et al. 2009).

Genomic analysis has revealed that this bacterium has a relatively large genome size, with approximately 4.6 to 5.6 million base pairs (Mbp), containing approximately 4,400 to 5,400 predicted protein-coding genes (Nikyri et al., 2012; Khayi et al., 2015; Zoledowska et al., 2018; Arizala and Arif, 2019). The *P. parmentieri* genome typically comprises a single circular chromosome, with few strains harboring a plasmid (Zoledowska et al., 2018). Numerous genes involved in important biological functions, including metabolism, replication, transcription, and translation have been found on the chromosome (Nikyri et al., 2012; Zoledowska et al., 2018). In addition, the genome contains many coding regions linked to pathogenicity, including those responsible for producing pectinolytic enzymes, extracellular enzymes, iron scavengers, toxins, protein effectors and specialized secretion systems (Nikyri et al., 2012; Zoledowska et al., 2018; Arizala and Arif, 2019). Together, these factors contribute to the infection and colonization of this pathogen inside its host plants.

Comparative genomic analyses have provided valuable insights into the genetic variations within the *P. parmentieri* species and their implications for host specificity and pathogenicity (Nykyri et al. 2012; Zoledowska et al. 2018a; Arizala & Arif 2019). Zoledowska et al. (2018a), reported several genes found to be part of the dispensable genome, indicating a high genomic plasticity across the *P. parmentieri* strains. These finding could be associated with the widespread distribution of this bacterium across the globe, its broad host range and its rapid adaptation to thrive in different climatic zones exhibiting different temperatures and humidity ranges (Zoledowska et al. 2018b). On the other hand, widely associated genome studies have shown that *P. parmetieri* has horizontally acquired gene regions crucial for pathogenicity and environment adaptation (Nykyri et al. 2012; Zoledowska et al. 2018a; Arizala & Arif 2019). The French and type strain RNS 08.42.1A (CFBP 8475^T^), possess several quorum sensing genes that seem to be acquired thorough horizontal gene transfer (Khayi et al. 2015). Moreover, the genome of the Finland strain SCC3193, which has been used as a model for studies during decades, was observed to harbor different genome islands, some of which contain important pathogenicity determinants. These outcomes provide a glimpse of the impact of gene transfer in the evolution of this phytopathogen.

No study has been conducted regarding the role of homologous recombination and its implication for the evolution of *P. parmentieri*. Considering this background, the primary objective of the present research article was to examine the importance of homologous recombination and its potential contribution to the evolutionary relationships, wide-strain genomic diversity, pathogenicity, host adaptation, and immunity of *P. parmentieri*. To our knowledge, this is a pioneering research that delves deeply into the study of homologous recombination across the core genome of the soft rot bacterium *P. parmentieri*.

## Results and Discussion

### Pan genome and population structural analyses reveal high diversity within strains of *P. parmentieri*

In total seventeen complete and fifteen draft genomes belonging to different *P. parmentieri* strains, isolated from various time periods and locations, were retrieved from the NCBI GenBank database and used in the analyses (**supplementary table S1**). The pan genome analysis conducted in Roary identified 3,742 and 71 genes in the core (genes present in 99% of strains) and soft core (genes present in 95%-99% of strains) genomes, respectively (**fig. 1A**). The accessory genome composed of the cloud and shell genomes, represented 55% of the pan-genome (8,433 genes). The shell genome (genes present in 15% - 94% of strains) and cloud genome (genes present in less than 15% of strains) accounted for 19% (1,605 genes) and 36% (3,015 genes) of the pan genome, respectively (**fig. 1A**). The higher number of genes identified in the accessory genome compared to the core/soft core genomes highlights the relevance of cloud and shell genes in the genetic diversity within the *P. parmentieri* species. Comparable results were found in a previous pan-genome study performed with 15 *P. parmentieri* strains, where 1,468 genes were part of the accessory genome and 1,847 were unique genes while the core genome was scarcely bigger than 50% of the pan-genome. Overall, these outcomes reflect an open pan-genome within *P. parmentieri,* which, as previously pointed (Zoledowska et al. 2018a), might be linked to the high genome plasticity and adaptation of this species to diverse environmental regions. Indeed, the presence of a large number of genes in the accessory genome (found also in our analysis) has been highlighted as the potential basis for the widespread diffusion of *P. parmentieri* (Zoledowska et al. 2018b) as the pathogen has been reported in numerous geographic locations worldwide, including, Finland, New Zealand, Canada, China, Turkey, South Africa and several states within the USA.

**Fig. 1.**
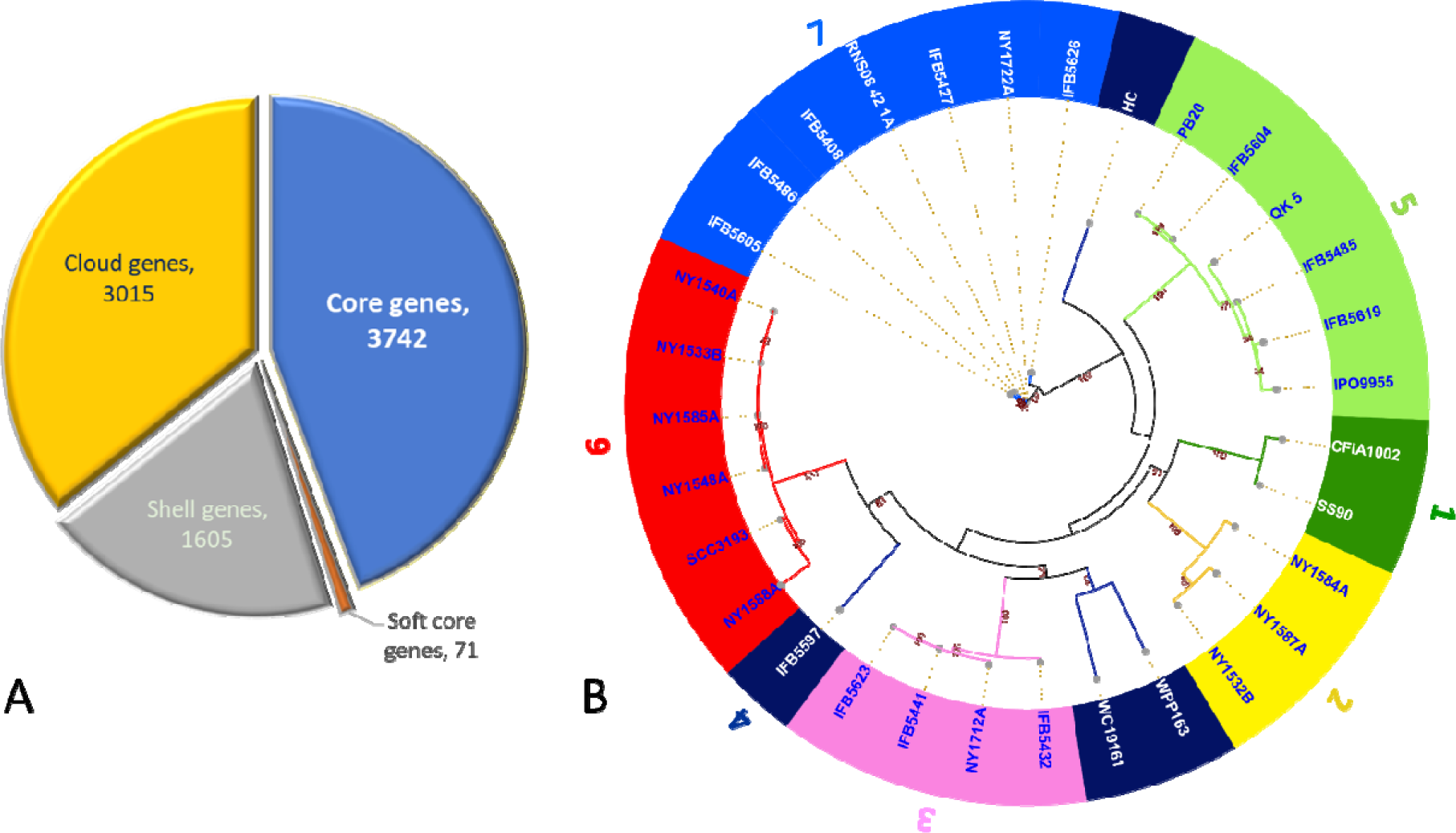
Pan-core genome and population structure analyses in *P. parmentieri*. (*A*) Pie chart displays the proportion of core, soft-core, shell and cloud genes in the 32 *P. parmentieri* genomes analyzed in the Roary pipeline. (*B*) Maximum likelihood tree depicting the population structure analysis of 32 *P. parmentieri* strains. The phylogenetic tree was built using iTOL v (https://itol.embl.de/) and the tips of each branch were annotated and colored according to the level clusters identified usin RhierBAPS and for matching the colors used by fastGEAR.

To examine the population structure within the 32 *P. parmentieri* strains, a clustering analysis was conducted using a hierarchical approach with Bayesian Analysis of Population Structure (BAPS, Cheng et al.,2013) implemented in R (RhierBAPS, Tonkin-Hill et al. 2018). RhierBAPS identified seven lineages (clusters) among the *P. parmentieri* strains (**fig. 1B**). The first lineage consisted of strains CFIA1002 and SS09 isolated in 2007 and 2017 in Canada: Alberta and Pakistan, Punjab, respectively. Lineage 2 encloses the New York (USA) strains NY1584A, NY1587A and NY1532B isolated in 2016. Lineage 3 comprises one New York (USA) strain NY1712A isolated in 2017 and three Poland strains, two collected in 2013 (IFB5441, IFB5432) and IFB5623 isolated in 2014. Interestingly, lineage 4 was the most heterogenous, composed of four isolates which formed like-outgroups subclades within the lineages 3, 5 and 6. Specifically, strains WPP163 and WC19161 isolated in 2004 (USA, Wisconsin) and 2019 (China, Hohhot), respectively, grouped separately but closely to the isolates assigned to the lineage 3, wherea IFB5597 isolated in Poland in 2014 grouped closely to the strains of lineage 6. The South Korea strain HC isolated in 2016 was distinctively separated from all lineages and clustered close to strains of lineage 5. Lineage 5, on the other hand, was integrated by six isolates, including two Poland strains isolated in 2014 (IFB5604, IFB5619), a Belgium strain (IFB5485) isolated in 2012, QK-5 isolated in 2019 in China (QingKou), the Russian strain PB20 isolated in 2014 in the Moscow region, and strain IPO1955 isolated in 2002 (unknown location). Like lineage 5, lineage 6 is constituted by six isolates, including five New York strains (NY1588A, NY1548A, NY1585A, NY1540A, NY1533B) isolated in 2016 and the Finland strain SCC3193 isolated in the early 1980s, used as a model for different molecular biology studies concerning *Pectobacterium* over many decades (Nykyri et al. 2012; Zoledowska et al. 2019). This strain was previously observed to cluster far away from the other *P. parmentieri* strains in a pan-genome tree due to the high number of unique genes present in this isolate (Zoledowska et al. 2018a). Here, we found that SCC3193 grouped together with strain NY1588A in a sub-clade within the lineage 6. Lastly, the lineage 7 harbors seven strains, including the type strain RNS 08-42-1ALJ isolated in France in 2008, four Poland strains isolated in 2013 (IFB5427, IFB5408) and 2014 (IFB5605, IFB5626), strain IFB5486 isolated from Belgium in 2012 and strain NY1722A isolated in New York in 2017. Except for strain IFB5427, which was isolated from weed, all other strains were isolated from either symptomatic tuber or potato stem.

Overall, all strains assigned to each of the seven lineages grouped differently regardless of their geographic origin and isolation year (**fig. 1B**), correlating with former phylogeographic analysis of this pathogen (Zoledowska et al. 2018b). This highly heterogenous pattern exhibited in the *P. parmentieri* population was suggested to be a consequence of high geographical mobility (Zoledowska et al. 2018b). Similarly, an elevated level of genetic diversity has been observed in the population structure analyses of the causal agent of tomato bacterial canker, *Clavibacter michiganensis,* as result of presumably multiple introductions events, where the international tomato seed trade has been highlighted as the leading force (Ansari et al. 2019; Thapa et al. 2017). Likewise, in *P. parmentieri*, the potato trading business appears to be one of the major contributors to the remarkably heterogeneity population of this pathogen and its global expansion.

### Reconstructed core-genome genealogy and rates of homologous recombination

To assess recombination events in the core genome of *P. parmentieri* and reconstruct its genealogy, a recombination analysis using the ClonalFrameML (Didelot & Wilson 2015) wa carried out employing a maximum likelihood tree as starter. ClonalFrameML detected 5,125 recombination events, with 3,094 occurring in the nodes that gave origin to the different lineage identified by RhierBAPS, and 2,031 accounted for recombination regions across the distinct *P parmentieri* strains used in this study (**fig. 2**). The ClonalFrameML algorithm estimated the following homologous recombination rates (**table 1**): the average length of recombined fragment (δ) was 83.456 bp, the average divergence between donor and recipient (ν) displayed a value of 0.0685 and the ratio of recombination to mutation rate was (*R/*θ) 0.236. This data shows that mutations have occurred 4.24 times more frequently compared to recombination. However, since each recombination event introduced an average of δν = 5.71 substitutions, the impact of recombination over mutation (*r/m*) was 1.348 times higher than mutation, emphasizing the relevance of recombination in *P. parmentieri* evolution.

**Fig. 2.**
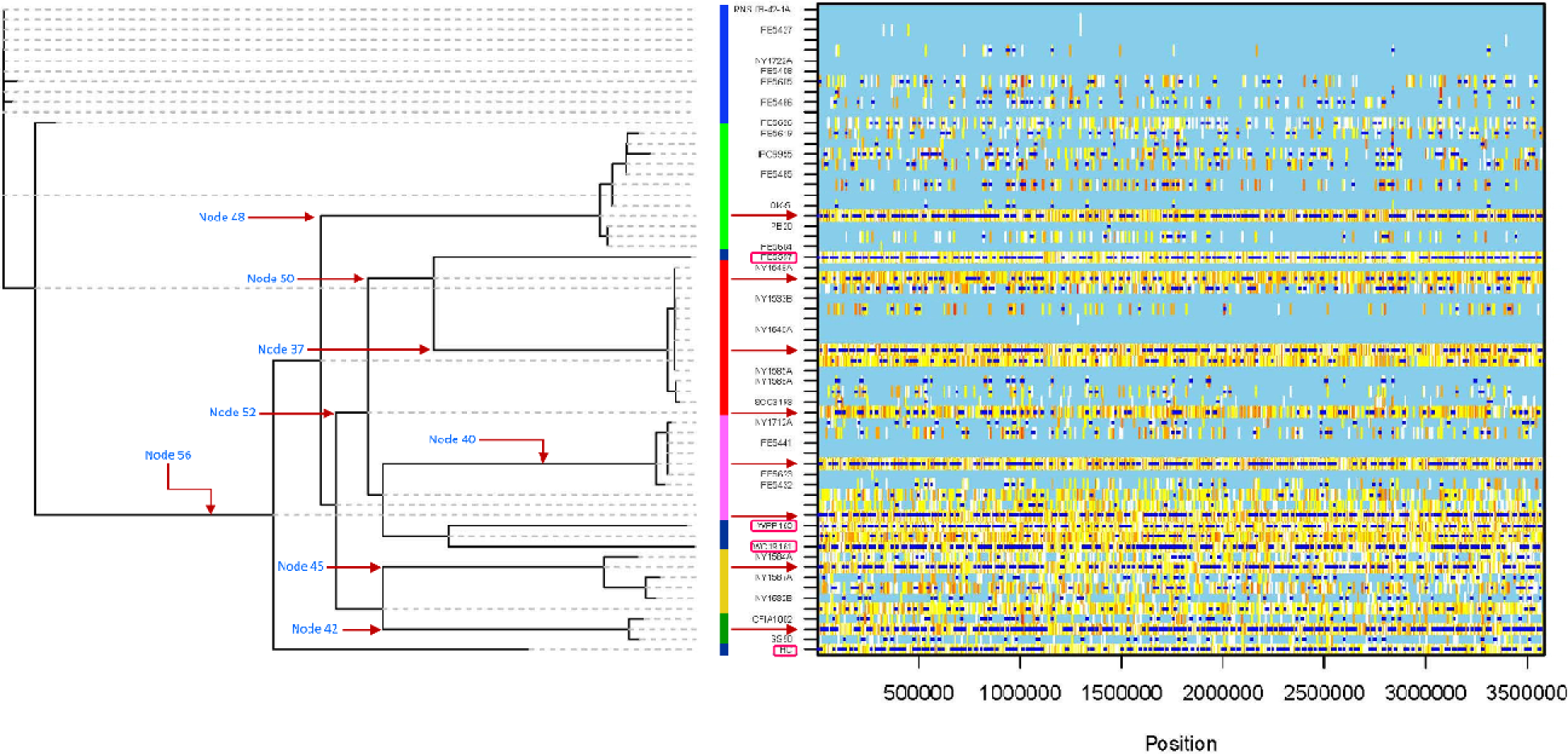
Reconstructed genealogy and inferred homologous recombination events in the core genome of 32 *P. parmentieri* strains are depicted in the graphic. The graphic depicts the phylogenomic relationships among the 32 isolates, along with locations of inferred recombination and substitutions events. Reconstructed maximum likelihood phylogenetic tree is depicted on the left; a color-coded vertical bar based on the lineages (clades) identified in RHierBAPS is shown in the center; inferred recombination events are illustrated in the right frame according to each position across the core genome (x-axis) and for each branch of the tree (y-axis). Unlabeled branches indicate ancestral lineages in the phylogeny. Dark blue horizontal lines pinpoint homologou recombination events, while vertical lines represent reconstructed substitutions or mutations. Light blue background color in th frame refers to no substitution or invariant polymorphic sites. White lines mark the absence of homoplasy, whereas the range of redness lines, from yellow to red, represent an increase of homoplasy levels. Inference of recombination analysis was conducted in ClonalFrameML using an RaxML tree with 1,000 bootstraps as a guide tree. Red arrows indicate the nodes with the highest recombination events, while pink framed squares point to the most recombinant strains as inferred by ClonalFrameML. Figure S1 shows the position and names of all nodes as assigned by ClonalFrameML.

**Table 1.**
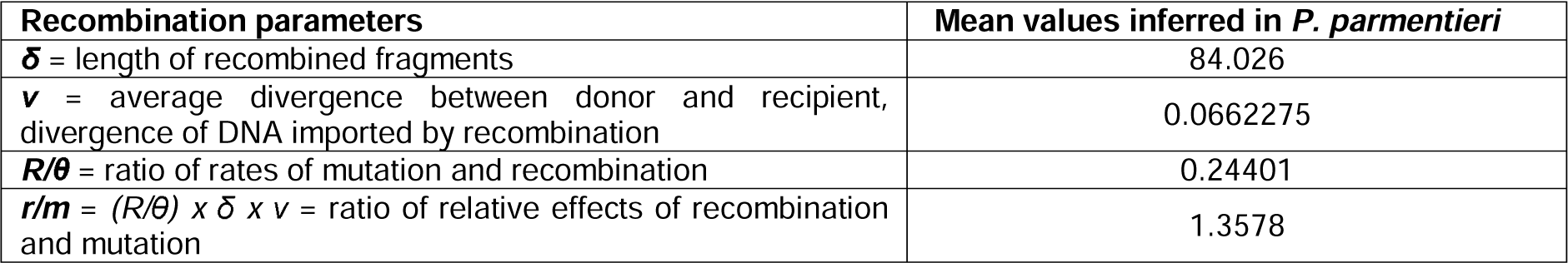
Recombination rates calculated in the core genome of *P. parmentieri* using ColnalFrameML.

In general, recombination events were detected in the ancestors of each lineage (**fig. 2**, indicated with a red arrow). **Supplementary figure S1** shows the positions and numbers of the nodes assigned by ClonalFrameML. The highest numbers of recombination events were observed in specific nodes, with node 48 having the most (343 events), followed by node 56 (341 events), node 42 (339 events), node 40 (338 events), node 37 (317 events), node 45 (292 events) and node 50 (150 events). Moreover, we observed distinct degrees of homoplasy (**fig. 2**, represented by yellow to red lines) across the lineages, seemingly imported as a consequence of mutation events occurring on the main branches of the genealogy tree. A similar effect has been reported in *Staphylococcus aureus* (Didelot & Wilson 2015). Intriguingly, all *P. parmentieri* strains assigned to the lineage 4 (**fig. 2**, marked in a pink square) were detected as the most recombinant isolates in the clonal phylogeny. The Poland strain IFB5597 led the list with 320 recombination events, featuring recombinant fragments ranging from 2,283 to 9 bp. The South Korean strain HC ranked second in the list, exhibiting 311 recombination regions with sizes ranging 1,995 to 3 bp. Following, the American strain WPP163, isolated in Wisconsin, presented 306 recombination events with fragment lengths ranging from 1,273 to 5 bp. Lastly, the Chinese strain WC19161 displayed 298 recombinant events with size ranging from 2,094 to 5 bp. Within the lineage 7, the Poland isolates IFB5626 and IFB5605 exhibited the highest number of recombinant events, with 72 and 60 inferred recombinant sites ranging from 1584 to 4 bp and 5,490 to 6 bp, respectively. Additionally, the Belgium strain IFB5486 showed 57 recombinant sites with a maximum and minimum lengths of 885 and 4 bp, respectively, whereas the type strain RNS 08-42-1ALJ, isolated from France, and the Poland strains IFB5408 and IFB5427, along with NY1722A from New York, appeared as non-recombinant strains. Regarding lineage 5, the strains IFB5619 (Poland) and IPO1955 (unknown origin) showed 70 and 95 recombination events, with size ranging from 935 to 6 bp and 1,335 to 8 bp, respectively. In contrast, the strains IFB5604 (from Poland) and QK-5 (from China) solely presented 1 and 10 recombination regions of 48 bp and 2,152 to 25 bp, respectively, while PB20 (from Russia) and IFB5604 (from Poland) did not show any recombination. In lineage 6, only two strains NY1588A and the model organism SCC3193, presented recombination events of 26 and 33, with fragments sizes oscillating between 5,490 to 88 bp and 1,016 to 6 bp, respectively.

On the other hand, the New York NY1712A and Poland IFB5432 strains clustering in lineage 3 displayed 36 and 29 recombinant regions with lengths ranging from 624 – 36 bp and 780 – 2 bp, respectively, while no recombination events were inferred in strains IFB5623 and IFB5441. All strains in lineage 2 have undergone through homologous recombination, thus recombination numbers of 68, 58, and 51 were observed in the New York strains NY1584A, NY1587A and NY1532B, with fragments sizes of 997 to 6 bp, 1,009 to 6 bp and 927 to 3 bp, respectively. In the case of lineage 1, the Canadian CFIA1002 (74 recombination events) and Pakistani strain SS90 (51 recombination events) strains showed recombinant sizes ranging from 1,261 to 4 bp and 606 to 5 bp, respectively. Altogether, these results indicate that the frequency of recombination varies across the strains. Related results were observed in recombination studies among *Xylella fastidiosa* subspecies (Vanhove et al. 2019). It seems reasonable that, similar to *X. fastidiosa* subspecies, due to geographic isolation, the introduction of foreign *P. parmentieri* isolates into new regions where other strains were already coexisting has contributed to homologous recombination events, leading to the emergence of novel strains.

### Screening of recombinant genes inferred in the *P. parmentieri* strains and ancestral nodes in the genealogy tree

All recombinant fragments predicted by ClonalFrameML in either each strain or the ancestral nodes in the genealogy tree were scanned and tracked using a customized BLASTN tool. Genes related with several vital cell functions, such as DNA repair, RNA methylation, RNA processing and degradation, pyrimidine and purine conversions, nitrate and nitrite ammonification, biosynthesis of amino acids (methionine, alanine, cysteine, glutamine, glutamate, aspartate, asparagine, lysine), utilization of D-galacturonate, D-glucoronate and D-ribose, uptake and utilization of lactose and galactose, phosphate metabolism, potassium homeostasis, biosynthesis of vitamins (thiamin, biotin, folate, pyridoxin), fatty acid biosynthesis, glycogen, glycerate and phosphate metabolism, protein chaperones and response to stress were detected in the analysis (**supplementary tables S2-S44**). Interestingly, recombination was also found to occur within the genes involved in the CRISPR-Cas system. The CRISPR-Cas system confers an innate and adaptive immunity in many bacteria by defending the bacterial cell against bacteriophages or other exogenous invaders (Koonin & Makarova 2013; Makarova et al. 2015). In the case of *P. parmentieri*, two types of CRISPR-Cas systems have been reported: subtype I-F and subtype I-E (Arizala & Arif 2019). In this research, recombination fragments were detected in the *cas* genes encoding the CRISPR-associated (Cas) proteins Cas1 (subtype I-F) and Cas3 (subtype I-E) with maximum sizes of 1,009 bp and 469 bp, respectively. Strains WPP163 and WC19161 exhibited recombination signals in both genes *cas1* and *cas3* (**supplementary tables S27-S28**). While strains HC and NY1587A showed recombination only in *cas1* (**supplementary tables S29, S38**). Conversely, in the strains IFB5597, NY1712A and NY1584A, homologous recombination events were detected solely in *cas3* (**supplementary tables S26, S36-S39**). Likewise, in the genealogy tree (**fig. 2**), recombination events within both *cas1* and *cas3* genes were also found in the ancestral nodes 37, 40 and 56, while nodes 41, 42, 48 and 53 showed recombination only in *cas3* gene, and node 45, and 50 exhibited recombination only in *cas1* (**supplementary tables S4-S22**). Altogether, this data suggests that homologous recombination within *cas1* and *cas3* derived from the main ancestral node 56 and was transmitted during evolution to the different lineage nodes, and finally, to the *P. parmentieri* strains mentioned above.

In line with our findings, previous studies have reported horizontal gene transfer (HGT) events in CRISPR-Cas regions (Chakraborty et al. 2010; Tyson & Banfield 2008). Arizala and Arif (2019) described the acquisition of an entire type III-A CRISPR-Cas system in *Pectobacterium aroidearum* PC1 from *Serratia* species through HGT mechanism. The transfer of *cas* genes between identical loci surrounding CRISPR regions has been reported in *Escherichia coli* and *Salmonella enterica* genomes, suggesting that CRISPR-associated regions might serve as hot spots for recombination of *cas* genes (Touchon & Rocha 2010). Additionally, homologous recombination has been implicated in facilitating this genetic exchange of *cas* genes (Touchon & Rocha 2010). Subsequently studies identified widespread microscale recombination events in several *cas* genes, highlighting the dynamic evolution of *cas* genes beyond *cas1* (Takeuchi et al. 2012). Building on these earlier investigations, the homologous recombination detected in *cas1* and *cas3* in *P. parmentieri* in our study support previous hypotheses about the coevolutionary competition of the CRISPR-Cas system as an immune defense mechanism against rapidly evolving viruses (Takeuchi et al. 2012; González-Torres et al. 2019).

Strikingly, recombination events were consistently appeared in genes integral to critical pathogenicity determinants, such as the type VI secretion system (T6SS), iron uptake systems (siderophore aerobactin, ferric citrate, enterobactin), lipopolysaccharides (LPS), plant cell wall degrading enzymes (CWDE), type IVb fimbrial low-molecular-weight protein/tight adherence protein (Flp/Tad) pilus, citrate metabolism, butanediol metabolism, flagella (**supplementary tables S2-S43**). These factors collectively play pivotal roles in the infection process, bacterial colonization, and the progression of soft rot/black leg diseases in *Pectobacterium* sp. (Arif et al. 2022; Arizala & Arif 2019; Nykyri et al. 2012; Charkowski 2018). This underscores the significant impact of homologous recombination on the evolution of several virulence-associated factors across the core genome of *P. parmentieri*.

The T6SS comprises 15-23 proteins, with 13 highly conserved proteins believed to be essential for its function (Shyntum et al. 2019). Although numerous functions such as virulence, bacterial competition and host interaction have been attributed to the T6SS (Morgado & Vicente 2022), a recent study has suggested its involvement in biofilm formation in *Pseudomonas aeruginosa* (Chen et al. 2020). Within our analysis, we found recombination regions in 10 T6SS-related genes (**supplementary tables S2-S28, S38**), including: *impA* encoding for the T6SS component TssA (ImpA) in the nodes 33 (252 bp) and 46 (98 bp), *impC* encoding for the T6SS component TssC (ImpC/VipB) in the strains IFB5597 (113 bp) and WC19161 (113 bp), *impE* encoding for the T6SS associated component TagJ (ImpE) in the node 49 (144 bp), *impF* encoding for the T6SS lysozyme-like component TssE in the nodes 40 (76 bp), 42 (261 bp), 45 (114 bp), 48 (375 bp), 56 (261 bp) and in the strains IFB5597 (364 bp), WPP163 (76 bp) and WC19161 (129 bp), *vasA* ecoding for the T6SS component TssF (ImpG/VasA) in the strain IFB5597 (30 bp), *vasC* encoding for the T6SS fork head associated domain protein ImpI/VasC in the node 44 (141 bp), *vasD* encoding for the T6SS secretion lipoprotein TssJ (VasD) in the node 37 (59 bp), v*asK* encoding for the T6SS component TssM (IcmF/VasK) in the node 44 (93 bp), *vasL* encoding for the Type VI secretion-related protein VasL in the nodes 46 (63 bp) and 37 (79 bp) as well as in the strain NY1587A (138 bp), *stk1* encoding for the T6SS Serine/threonine protein kinase PpkA in the node 52 (104 bp). Additionally, ClonalFrameML detected recombination events in the genes encoding for the products haemolysin-coregulated protein (Hcp) in the nodes 40 (306 bp) and 43 (48 bp), in the strains HC (12 bp) and SS90 (93 bp), and the valine-glycine repeat protein (VgrG) in the nodes 36 (220 bp) and 40 (220 bp) (**supplementary tables S3-S9, S29, S33**). Both Hcp and VgrG are considered as major effector proteins injected from T6SS into host cells (Pukatzki et al. 2006; Mattinen et al. 2007). Importantly, the ImpC protein, a T6SS element showing recombination in our analysis, has been identified as a crucial component of the T6SS, playing a role in virulence and participating in the intracellular host response in *Pseudomonas syringae* pv. *tomato* (Sarris et al. 2010). Moreover, T6SS gene clusters are often found within predicted genomic islands, suggesting the potential of this system for horizontal gene transfer between bacteria (Morgado & Vicente 2022). Nykyri et al. (2012) reported two T6SS in the genome of the Finland strain *P. parmentieri* SCC3193, with one located in a genomic island. Our findings further underscore that in *P. parmentieri*, homologous recombination seems to be another important mechanism in the evolution of the T6SS core genes. Consistent with our findings, frequent homologous recombination has been observed in *Vibrio cholera* and is considered a major contributor to the high diversity of T6SS effector genes (Kirchberger et al. 2017). Comprehensive studies are warranted to establish a conclusive understanding of the specific role of homologous recombination in the evolution of the T6SS in *P. parmentieri*.

Interestingly, ClonalFrameML inferred recombination events in three different lipopolysaccharides (LPS) encoding gene clusters. Polysaccharides are widely recognized bacterial virulence factors, serving various functions, including attachment inside the cell host, protection from plant toxins and adverse environmental conditions, and favoring host colonization (Toth et al. 2006). Concerning the enterobacterial common antigen (ECA), recombination was predicted in five genes: *wzxE* (lipid III flippase) found in the nodes 36 (647 bp), 37 (145 bp), 39 (647 bp), 44 (1050 bp), 46 (1115 bp), 52 (2152 bp), 57 (1238 bp), 58 (2152 bp), 61 (1245 bp) and in the strains IFB5597 (345 bp), WPP163 (345 bp), WC19161 (2094 bp), SCC3193 (512 bp), QK-5 (2152 bp), NY1588A (2162 bp) and IFB5432 (445), *rffG* (dTDP-glucose 4,6-dehydratase) found in the nodes 37 (309 bp), 39 (422 bp), 46 (524 bp), 52 (524 bp), 57 (522 bp), 58 (522 bp), 61 (357 bp) and in the strains IFB5597, WPP163, WC19161, NY1588A and IFB5432, *wecC* (UDP-N-acetyl-D-mannosamine dehydrogenase) and *wecA* (undecaprenyl-phosphate alpha-N-acetylglucosaminyl 1-phosphate transferase) found merely in the nodes 56 (110 bp) and 37 (65 bp), respectively (**supplementary tables S3-S43**). The ECA plays a major role in bacterial physiology, influencing the host immune response and participating in host-pathogen interactions (Rai & Mitchell 2020). Beyond ECA, the exopolysaccharide (EPS) and O-antigen cluster also underwent recombination in two genes (**supplementary tables S18-S37**): *gnd* (6-phosphogluconate dehydrogenase, decarboxylating) with recombined fragments observed in the nodes 52 (504 bp), 57 (465 bp), 58 (504 bp), 61 (183 bp) and in the isolates IFB5597 (336 bp), WC19161 (715 bp), IFB5626 (1335 bp), IPO1955 (1335 bp), QK-5 (321 bp) and NY1588A (1378 bp), and *wza* (putative polysaccharide export protein YccZ precursor) found in the nodes 52 (1129 bp) and 58 (1094 bp) and in the strains WC19161 (1079 bp), QK-5 (1094 bp), NY1588A (1100 bp). The lipo-oligo/polysaccharide cluster (LOS/LPS) presented recombination in three genes (**supplementary tables S4-S32**): *waaC* (lipopolysaccharide core heptosyltransferase I) found in the nodes 37 (411 bp), 40 (238 bp), 42 (505 bp), 43 (116 bp), 46 (799 bp), 50 (117 bp), 53 (105 bp), 56 (372 bp) and in the strains IFB5597 (233 bp), WC19161 (115 bp), IPO1955 (505 bp) and IFB5619 (435 bp), *rfaD* (ADP-L-glycero-D-manno-heptose-6-epimerase) found in two nodes 37 (96 bp) and 50 (81 bp), and *waaL1* (O-antigen ligase) found only in the node 43 (196 bp). Our results show that homologous recombination actively contributes to the evolution of polysaccharide clusters, facilitating genetic exchange within the *P. parmentieri* population. Consistent with our findings, recombinant alleles in LPS genes have been described in evolutionary studies of plant pathogens such as *Xylella fastidiosa*, *Xanthomonas euvesicatoria,* and *X. perforans* (Jibrin et al. 2018; Potnis et al. 2019). Moreover, we speculate that the observed recombination homologous in different LPS-genes might be involved with the recently described heterogeneity within the LPS structure of certain *P. parmentieri* strains (Ossowska et al. 2022).

The ability to acquire iron constitutes a relevant virulence determinant during bacterial pathogenesis, especially in iron-deficient environments due to metal ion restriction adopted by the host as a nutritional immunity strategy (Palmer & Skaar 2016; Mosbahi et al. 2018). Recombination events were predicted in genetic regions associated with three different iron uptake systems: the ferric citrate (*fecIRABCDE*) transport system, the siderophore aerobactin (*iuc*) and the siderophore enterobactin (*ent*). In the ferric citrate cluster (**supplementary tables S4, S6-S16, S39**), the gene *fecD*, encoding for the iron (III) dicitrate transport system permease protein FecD, manifested recombination in the nodes 37 (33 bp), 40 (78 bp), 41 (684 bp), 42 (54 bp), 48 (342 bp), 50 (209 bp) and only in the isolate NY1584A (54 bp). In pathogenic Gram-negative bacteria, this system is formed inside plant tissues and enabling long-distance iron transport (Franza & Expert 2013; Gorshkov et al. 2021). On the other hand, the siderophore aerobactin presented signals of recombination in four genes (**supplementary tables S4-S43**): *fhuB* (ferric hydroxamate ABC transporter, permease component FhuB) showing recombination in the nodes 48 (566 bp) and 50 (397 bp), and in the strains IFB5597 (295 bp), WPP163 (806 bp), WC19161 (62 bp), HC (62 bp), IFB5626 (62 bp) and SS90 (29 bp), *fhuC* (ferric hydroxamate ABC transporter, ATP-binding protein FhuC) found solely in IPO1955 (147 bp), and *iucC* (aerobactin biosynthesis protein IucC) found in the nodes 54 (135 bp) and 57 (314 bp) and strain IFB5432 (181 bp), and the iron-chelator utilization protein with recombination regions observed in the nodes 37 (81 bp), 40 (31 bp) and 48 (31 bp), and the isolates WPP163 (211 bp) and NY1584A (178 bp). Importantly, Ling et al. (2013) reported that the aerobactin synthesis gene, *iucC,* plays a significant role in the pathogenicity of *Escherichia coli* O2 strain E058. Recombination in the siderophore enterobactin was only detected in the node 56 (288 bp) in the gene encoding a transcriptional regulator of the AraC family, enterobactin-dependent (**supplementary table S22**). In *P. atrosepticum*, this siderophore has been hypothesized to confer resistance to different stressors and contribute to virulence (Gorshkov et al. 2021). Additionally, a coding sequence annotated as ferric iron ABC transporter, permease protein, and catalogued in the subsystem “iron acquisition in *Streptococcus*” displayed recombination events in six ancestral nodes (**supplementary tables S4-S42**): 37 (183 bp), 42 (643 bp), 48 (62 bp), 49 (123 bp), 53 (91 bp), 56 (123 bp), and six isolates: IFB5597 (91 bp), WPP163 (408 bp), HC (148 bp), NY1712A (48 bp), NY1532B (121 bp), IFB5486 (221 bp). In general, these distinct recombination rates observed in different iron-scavenging genes might reflect an adaptative mechanism adopted by *P. parmentieri* to successfully colonize the plant tissue and survive in poorly-iron environments.

The secretion of cell wall degrading enzymes (CWDE) is the most well-recognized and the most studied pathogenicity determinant in pectinolytic bacteria of the genera *Pectobacterium* and *Dickeya*; indeed, efficient production of CWDE is directly linked with soft rot and black leg symptoms (Nykyri et al. 2012; Charkowski 2018; Arizala & Arif 2019; Boluk et al. 2021; Arif et al. 2022). In our study, recombining signatures were detected mainly in the *pelX* gene (**supplementary tables S6, S11, S26-S29**), encoding for an exopolygalacturonate lyase, in the strains IFB5597 (267 bp), WC19161 (225 bp) and HC (91 bp) as well as in the ancestral nodes 40 and 45 (91 bp). The pectin methylesterase, *pemB*, also exhibited recombination in the node 49 (44 bp; **supplementary table S15**), while in the case of the oligogalacturonides degradation, the *kdfF* (pectin degradation protein KdgF) locus underwent recombination in the Wisconsin strain WPP163 (81 bp; **supplementary table S27**) and in the ancestral nodes 42 (81 bp; **supplementary table S8**) and 45 (91 bp; **supplementary table S11**). In agreement with our analysis, recombination regions have also been encountered in the CWDE-genes of the pathogen *Xylella fastidiosa* (Potnis et al. 2019). We also observed recombination in genes encoding for proteases and peptidases. For instance, a recombinant fragment of 98 bp was identified in the gene *pqqL* (putative zinc protease) in the node 51 (**supplementary table S17**). On the other hand, three loci (*citC*, *dcuC*, *citF*) involved with citrate metabolism, uptake and regulation showed evidence of recombination. The genes *citC* (citrate [pro-3S]-lyase ligase) and *citF* (citrate lyase alpha chain) displayed recombination fragments of 78 bp and 321 bp, only in the strain HC and node 48, respectively (**supplementary tables S14, S29**). Conversely, *dcuC* has undergone recombination in four strains (SCC3193, NY1588A, IFB5605 and IFB5432; **supplementary tables S34, S37, S41, S43**) and four nodes (33, 46, 49 and 57; **supplementary tables S2, S12, S15, S23**). The New York strain NY1588A harbored the largest recombined fragment of 1,263 bp (**supplementary table S37**). The citrate uptake system was reported to be critical for bacterial virulence of *P. atrosepticum* by reducing the citrate concentration during colonization in potato tubers (Urbany & Neuhaus 2008). Our findings highlight the relevance of homologous recombination in the evolution of the citrate uptake system and its potential association with the adaptation of certain *P. parmentieri* strains to host environments containing high citrate levels. The 3-hydroxy-2-butanone (3H2B – acetoin, butanediol) pathway has been characterized as another relevant factor for *P. carotovorum* pathogenesis by promoting media alkalization and, consequently, favoring plant cell maceration (Marquez-Villavicencio et al. 2011). In our analysis, the strains IFB5597 (36 bp), WC19161 (60 bp) and ancestral nodes 40 (147 bp) and 45 (60 bp) experienced recombination events in the gene product acetolactate synthase large subunit (**supplementary table S6, S11, S26-S28**).

The Flp/Tad (Fimbrial low-molecular-weight protein/Tight adherence protein) pilus, encoded by the *flp/tad* genes, was characterized as another novel virulence determinant in *Pectobacterium* due to its implication with maceration of potato tubers (Nykyri et al. 2013). ClonalFramelML revealed recombination events in four genes of the *flp/tad* biogenesis cluster (**supplementary tables S2-S8, S16-S31**). The *tadA* (type II/IV secretion system ATP hydrolase TadA/VirB11/CpaF, TadA subfamily) gene showed recombination in the node 37 (96 bp), 42 (66 bp), 57 (79 bp) and strain WC19161 (102 bp). Recombination in the *rcpC* (Flp pilus assembly protein RcpC/CpaB) gene was found in the nodes 40 (42 bp), 50 (52 bp) and in the strains HC (85 bp) and IF5626 (52 bp). The *tadC* (type II/IV secretion system protein TadC, associated with Flp pilus assembly) gene displayed recombination regions in the nodes 33 (21 bp) and 52 (21 bp), whereas recombination within the *rcpB* (Flp pilus assembly protein RcpB/CpaD) locus was predicted only in the node 57 (32 bp). A comparative genomics study showed that the *flp/tad* cluster is harbored by all *Pectobacterium* species (Arizala & Arif 2019). The discovery of recombination signals in some of the *flp/tad* genes highlights how this system is evolving via genetic exchange in *P. parmentieri*, presumably linked to the host adaptation of the pathogen to efficiently macerate tissues.

The flagella-encoding cluster has shown to be pivotal for motility, colonization, and virulent lifestyle in numerous pathogenic bacteria, including *Pectobacterium* (Toth et al. 2006; Arizala & Arif 2019; Nykyri et al. 2012). Three flagella-genes have undergone recombination, including *fliD*, *fliK*, *flgI* (**supplementary tables S14-S16, S26-S29, S41**). While recombinant fragments of 155 bp located in node 50 and 300 bp in strain IFB5605 were found in the loci *fliD* (flagellar cap protein FliD) and *fliK* (flagellar hook-length control protein FliK), respectively, the *flgI* (flagellar P-ring protein FlgI) gene displayed recombination in four strains (IFB5597 – 286 bp, WPP163 – 286 bp, WC19161 – 393 bp and HC – 387 bp) and two nodes (48 – 279 bp and 50 – 99 bp). In a former study, the flagellar genes were shown to participate in chemotaxis, motility and were fundamental for pathogenesis of *D. dadantii* (Jahn et al. 2008). Moreover, a reduced virulence was observed in tobacco plants infected with *P. parmentieri* SCC3193 mutants lacking motility (Pirhonen et al. 1991). In another report, the flagellum was related with the secretion of colin, a toxin released to kill competitors in the niche, thereby suggesting an antagonist role of the flagellar apparatus in *Pectobacterium* (Charkowski et al. 2012). Lastly, the gene predicted as *pgaC,* encoding for the biofilm PGA synthesis N-glycosyltransferase PgaC protein, showed a recombination region of 246 bp in the strain WC19161. PgaC was demonstrated to participate in host-bacteria interactions and modulate biofilm formation in the human pathogen *Klebsiella pneumoniae* (Chen et al. 2014).

### Insights into ancient and recent recombination events in *P. parmentieri*

To examine the ancestral and recent recombinogenic regions between and within the seven lineages (inferred by RhierBAPS – **fig. 1B**), we used the fastGEAR software (Mostowy et al. 2017), utilizing the core-genome alignment obtained from the Roary pipeline as input data. Ancestral recombinations are defined as recombinant regions present in all strains belonging to the lineage (Mostowy et al. 2017). Since the lineage 7 contained the largest number of strains within the *P. parmentieri* population, according to the principle of parsimony, this lineage was assigned as the main donor, while the other lineages (1-6) served as recipients for the ancestral recombination analysis. Thus, excluding lineage 7, all other lineages have undergone recombination, as depicted by **figure 3A**, where a mosaic like pattern across the core genome is evident. In total, 486 ancient recombination events were detected in the core genome obtained from the 32 *P. parmentieri* isolates (**fig. 3B**).

**Fig. 3.**
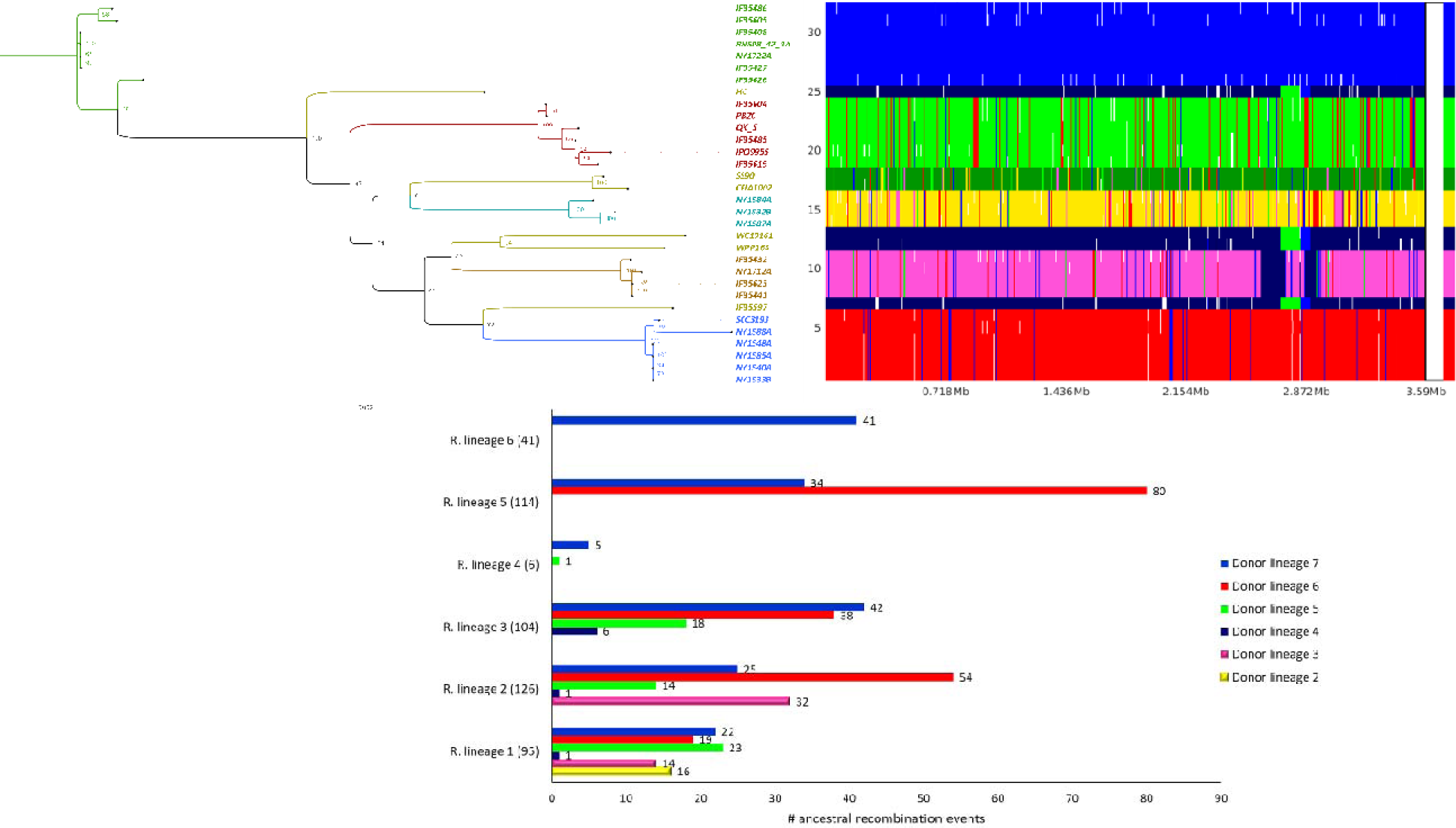
Ancestral recombination events identified using fastGEAR across the core genome of 32 *P. parmentieri* strains are illustrated. (*A*) Maximum likelihood phylogeny, using a bootstrap support of 1,000 replicates, is illustrated on the left, whereas panel showing the recombination sites and the population genetic structure is located on the right. The y-axis on the panel displays the 32 strains present in the core genome alignment, and the x-axis displays the sequence positions. Strains on the y-axis wer positioned according to the ML phylogeny. Seven lineages previously defined with RhierBAPS were used in the analysis. Th vertical color-coded bar on the right side of the panel shows the division of the 32 strains into the seven predicted lineages. Each lineage represents a specific clade and is labelled and colored as indicated in the **supplementary figure S2**. The black color denotes recombination from outside origin of any lineage in the data set. The presence of different color lines within a lineage in the frame pinpoints ancestral recombination event sites occurred respect to the sequence position. White gaps in the panel show the recent recombination events that were removed before the analysis of ancestral recombination. (*B*) Horizontal bar graphic illustrating the number of ancestral recombinations that were received in each lineage. The bars were colored based on the color of the lineages previously assigned in figure 1B.

The overall size of ancestral recombination events ranged 43 bp to 119 kb, with a median length of 4.8 kb. Lineage 2 exhibited the highest number of recombinations, totaling 126 events originating from all six lineages (lineage 2 – 16, lineage 3 – 14, lineage 4 – 1, lineage 5 – 23, lineage 6 – 19 and lineage 7 – 22). This finding highlights the three New York strains (NY1584A, NY1532B, NY1587A) belonging to the lineage 2 as highly ancient recombinants. In contrast, lineage 5 presented 114 ancestral recombination events, with fragments donated by lineages 6 (80 events) and 7 (34 events). Notably, this lineage transferred the highest number of recombinogenic regions (191 events) to the other lineages, potentially resulting from coexistence of this lineage with the other ones in the same ecological niche, promoting DNA recombination. In *Xyllela fastidiosa*, for instance, homologous recombination was reported in the South American strains sharing the same geographic area (Coletta-Filho et al. 2017). Lineage 4 has experienced the lowest level of ancestral recombination events, with 1 and 5 fragments donated by lineages 5 and 4, respectively. In the case of lineage 6, 41 events were detected, originated from lineage 7.

Regarding recent recombinations, these events refer to recombinant regions present in a subset of strains within a lineage (Mostowy et al. 2017). Similar to the ancestral recombination plot, a mosaic like-pattern indicative of recent recombination signals was observed in the core genome of most of the strains (**fig. 4A**). A total of 941 unique events were detected, with some events repeating across the strains, resulting in a total of 1,177 events (**fig. 4B**). Upon scanning these recent recombination regions, we identified 412 genes that have undergone recent recombination, which represents 11 % of the *P. parmentieri* core genome. The recombinant fragments displayed an average length of 1.38 kb, ranging from 36 bp to 13.96 kb. The twofold number of recent recombination events compared to the ancestral ones highlights that *P. parmentieri* is continuously evolving and adapting to new environments. This observation aligns with the widespread distribution of this bacterium reported in many countries in recent years (Pitman et al. 2010; Pasanen et al. 2013; Zoledowska et al. 2018b; Arizala et al. 2019). No signs of homologous recombination could be inferred in the strains IFB5408, RNS 08-42-1A, NY1722A, IFB5427, IFB5623 and IFB5441. In contrast, the strains IFB5626, IFB5597, HC and NY1584A showed high levels of recombinants with 140, 122, 113 and 87 events, respectively (**fig. 4B**). Strains CFIA1002, QK-5 and SS90 experienced a relatively low rate of recombination with 9, 10 and 13 events, respectively. Lineage 1 appeared as the major donor with 272 events, donating mainly to the strains IPO 9965 (24%), IFB5626 (21%) and NY1584A (18%). On the contrary, 5% of the total recent recombination events were categorized as outside origins and identified in 10 strains: HC, IFB5485, IFB5486, IFB5597, IFB5626, NY1588A, PB20, SCC3193, WC19161 and WPP163. These outside origin events are defined as recombined regions originating from outside the population or not corresponding to any of the lineages found in the population (Mostowy et al. 2017). Strains NY1588A and IFB5597 led the highest position in this category, each exhibited 15 and 16 outside origin events, respectively. Previous studies have documented the coexistence of soft rot bacteria from the genera *Pectobacterium* and *Dickeya* in the same niche (Ge et al. 2021; Degefu 2021; Kastelein et al. 2021). Considering this, it seems logical that these external origins regions might have been incorporated in the genomes of these 9 *P parmentieri* strains through genetic exchange with either other *Pectobacterium* species or members of the *Dickeya* genus.

**Figure 4.**
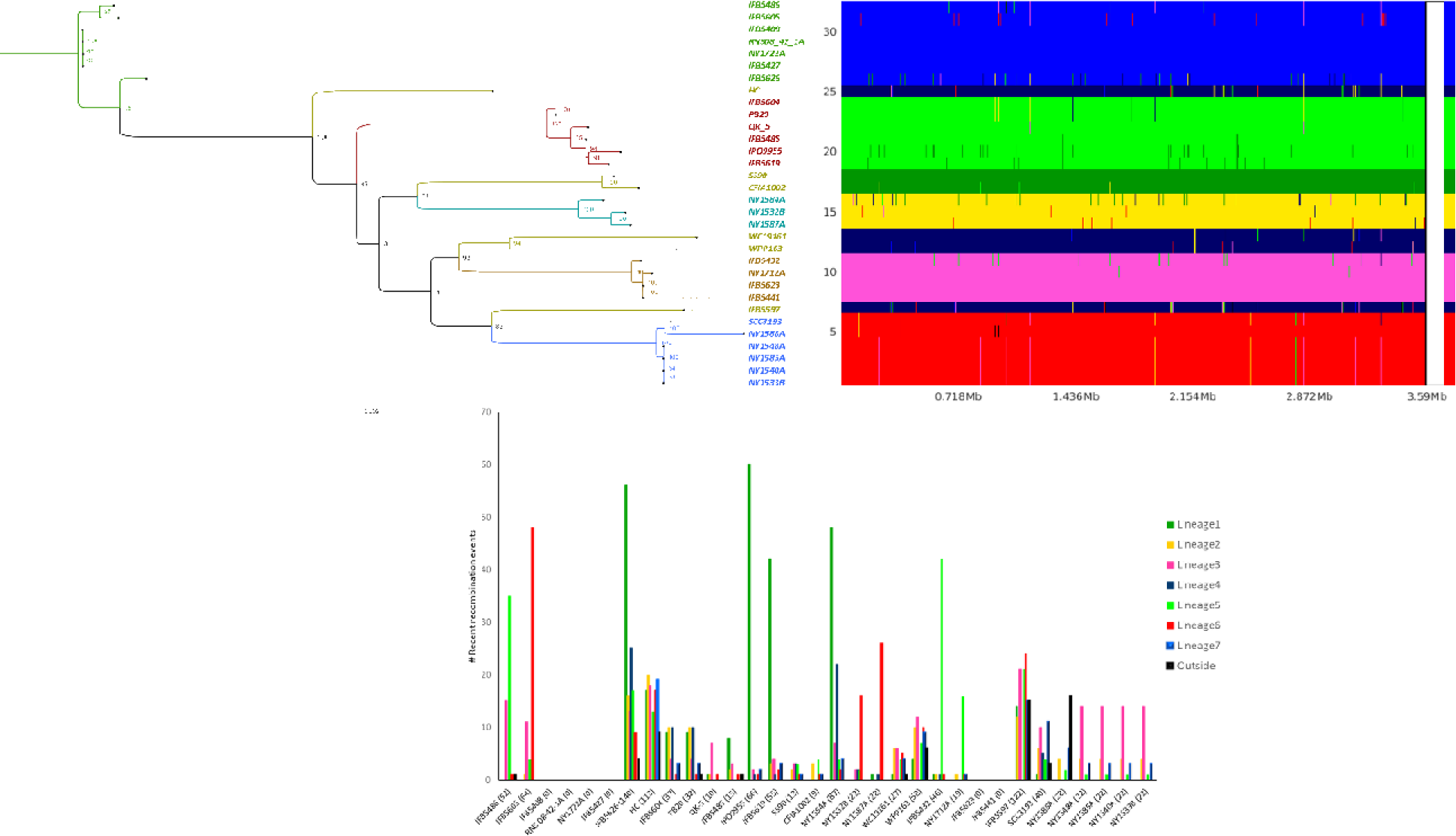
Recent recombination events identified using fastGEAR across the core genome of 32 *P. parmentieri* strains ar illustrated. (*A*) The y-axis on the panel displays the 32 strains present in the core genome alignment, and the x-axis displays the sequence positions. Strains on the y-axis were positioned according to the ML phylogeny depicted on the left. Seven lineages previously defined with RhierBAPS were used in the population clustering analysis. The vertical color-coded bar on the right side of the panel shows the division of the 32 strains into the seven predicted lineages. Each lineage represents a specific clade and is labelled and colored as indicated in the **supplementary figure S1**. Black color denotes recombination from outside origin of any lineage in the data set. The presence of different color lines within a lineage in the frame pinpoints recent recombination event sites occurred respect to the sequence position. (*B*) Bar graphic depicting the number of recent recombinations received by each strain. The bars were color-coded based on the color of the lineages assigned in **supplementary figure S2**. Maximum likelihood tree based on the core genome alignment of 32 *P. parmentieri* strains. The positions and names of all nodes assigned by ClonalFrameML are indicated and highlighted with different colors. The tree was visualized using FigTree v1.4.4 and mid-point rooted.

Subsequently, we traced and identified the coordinates and functional implications of all gene regions affected by ancestral (**supplementary tables S45-S50**) and recent recombination events (**supplementary tables S51-S76**). Genes associated with cellular functions, including DNA repair, DNA replication, RNA methylation, RNA processing and degradation, tRNA aminoacylation, cell respiratory pathways, dehydrogenase complexes, purine and pyrimidine conversions, fatty acid biosynthesis, redox cycle, lactose and galactose utilization, sucrose utilization, glycogen metabolism and oxidative stress have been affected by both ancestral and recent recombination events (**tables 2-3**). This underscores the significance of homologous recombination in the evolution of *P. parmentieri*. Recombinant fragments from recent and ancestral events were also observed in genes encoding amino acids and vitamins biosynthesis, phosphate metabolism, copper and potassium homeostasis, nitrite and nitrate ammonification, carbon starvation, nitrosative stress and ammonia assimilation (**tables 2-3**). In *X. fastidiosa* and *Xanthomonas* species, recombination in these loci has been associated with an adaptability mechanism for surviving in niches with limited nutrient sources (Chen et al. 2018; Potnis et al. 2019).

**Table 2.**
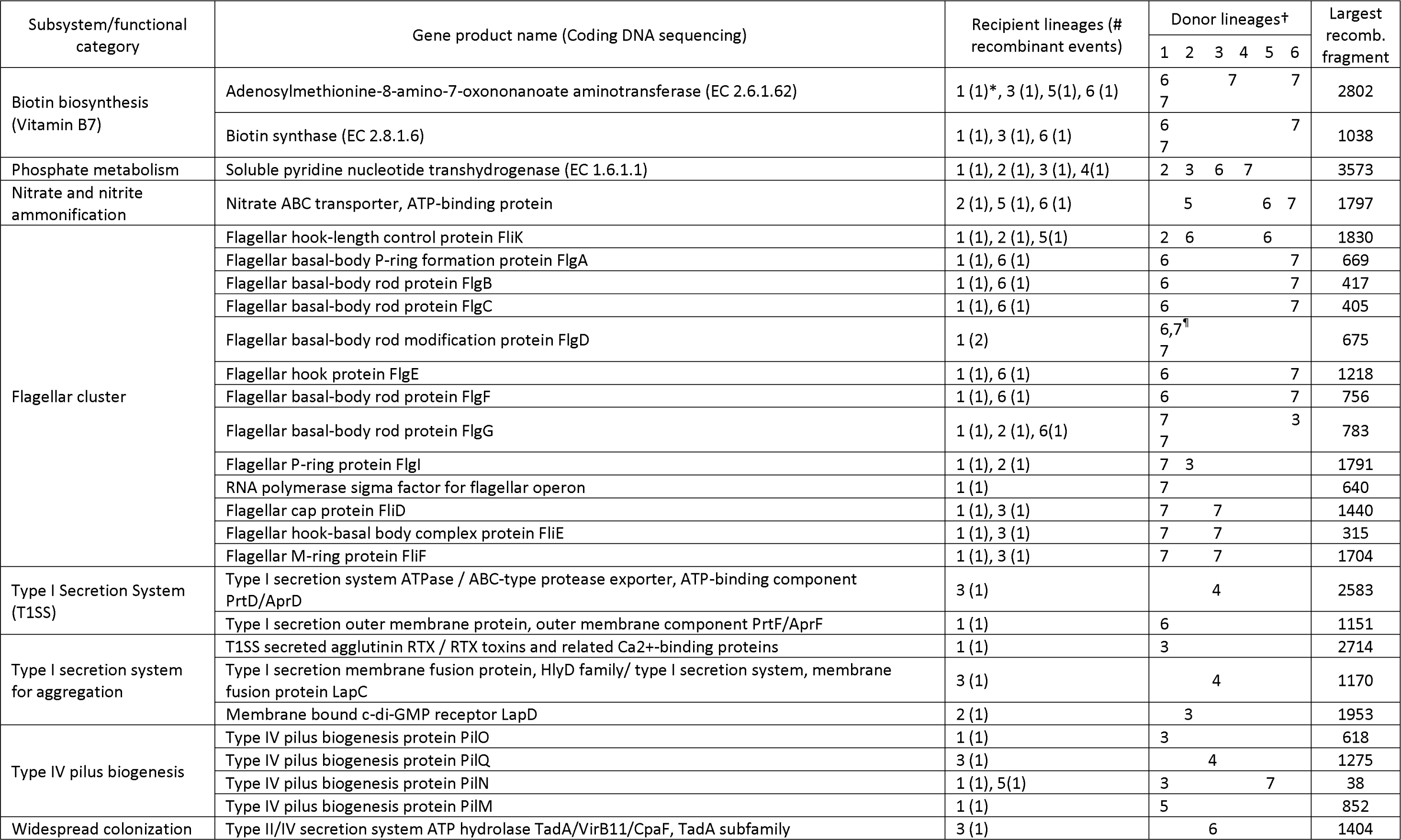
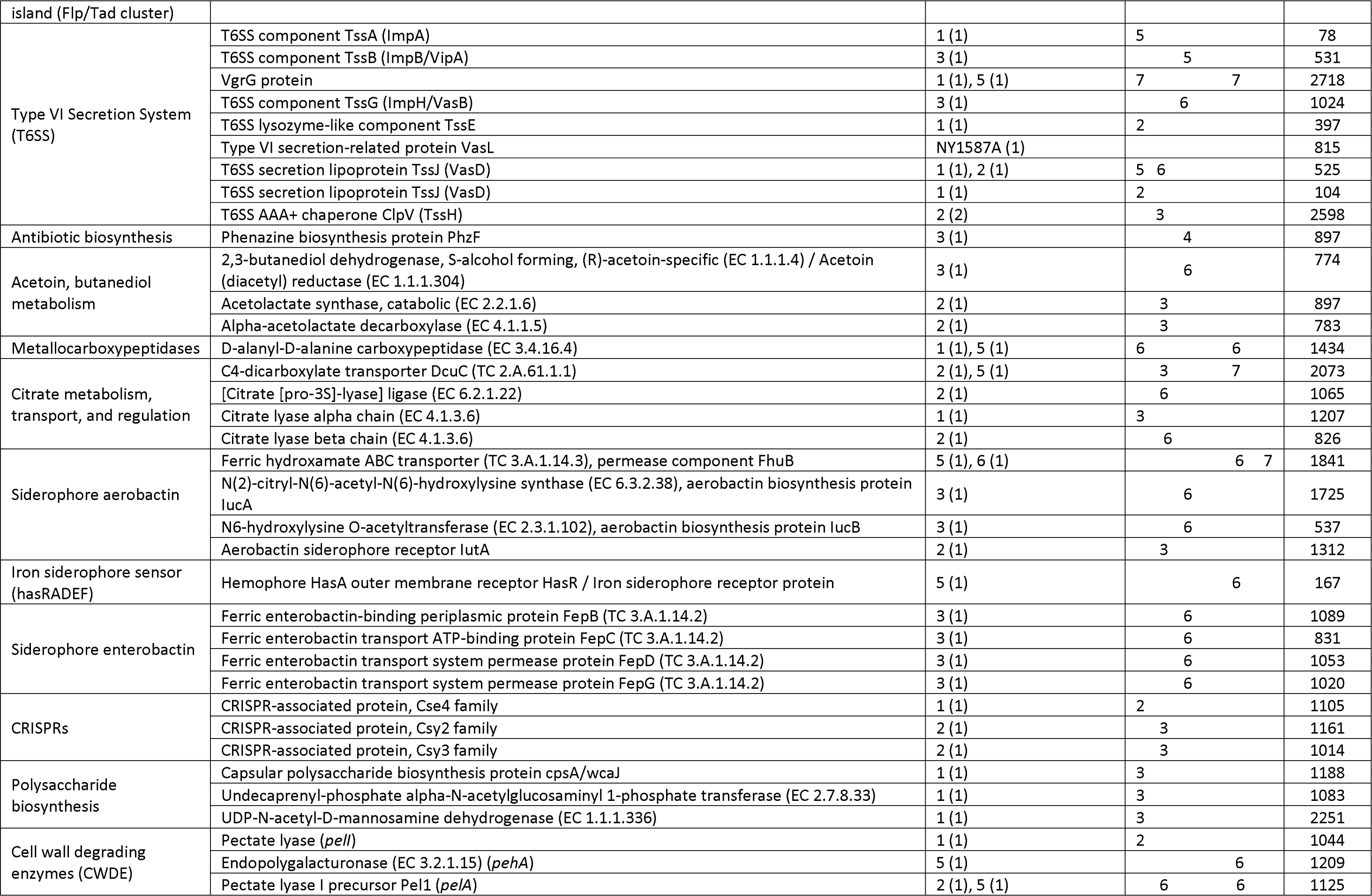
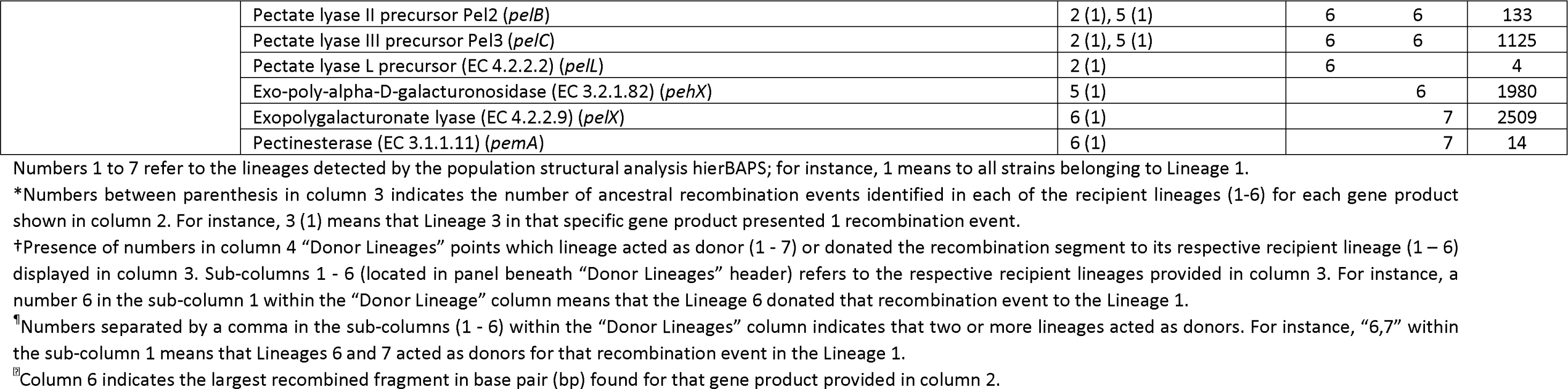
Important ancestral recombination events predicted in the core-genome of *Pectobacterium parmentieri*. The genes where the recombination events were identified are listed per subsystem and their product name is described as annotated in the RASTtk server.

**Table 3.**
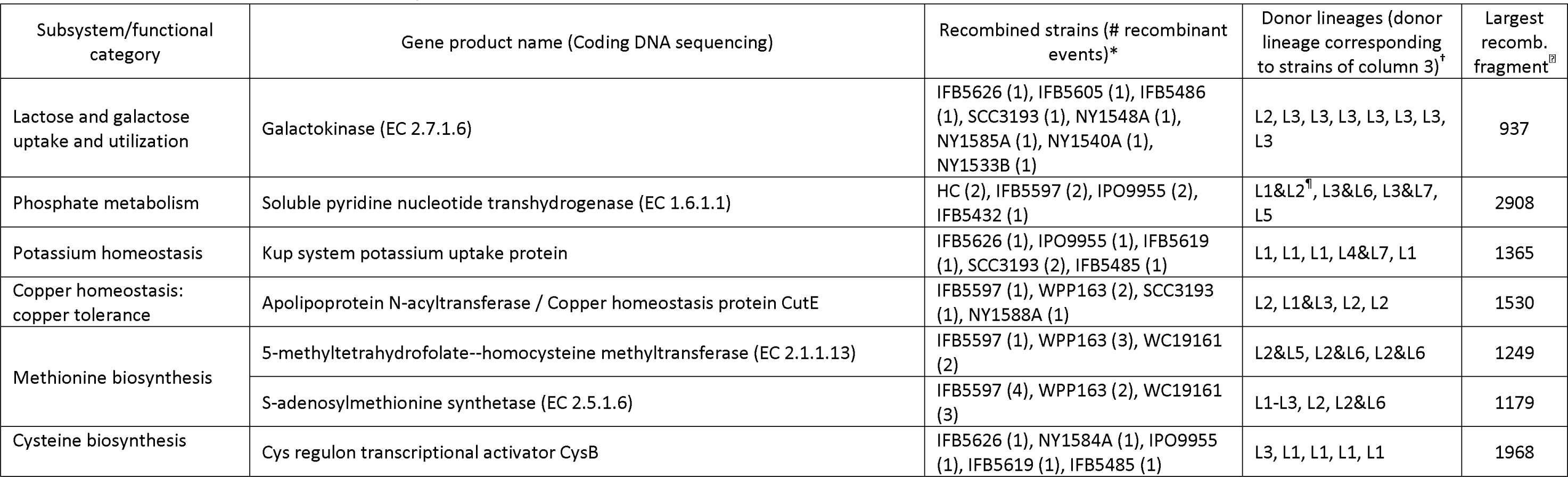
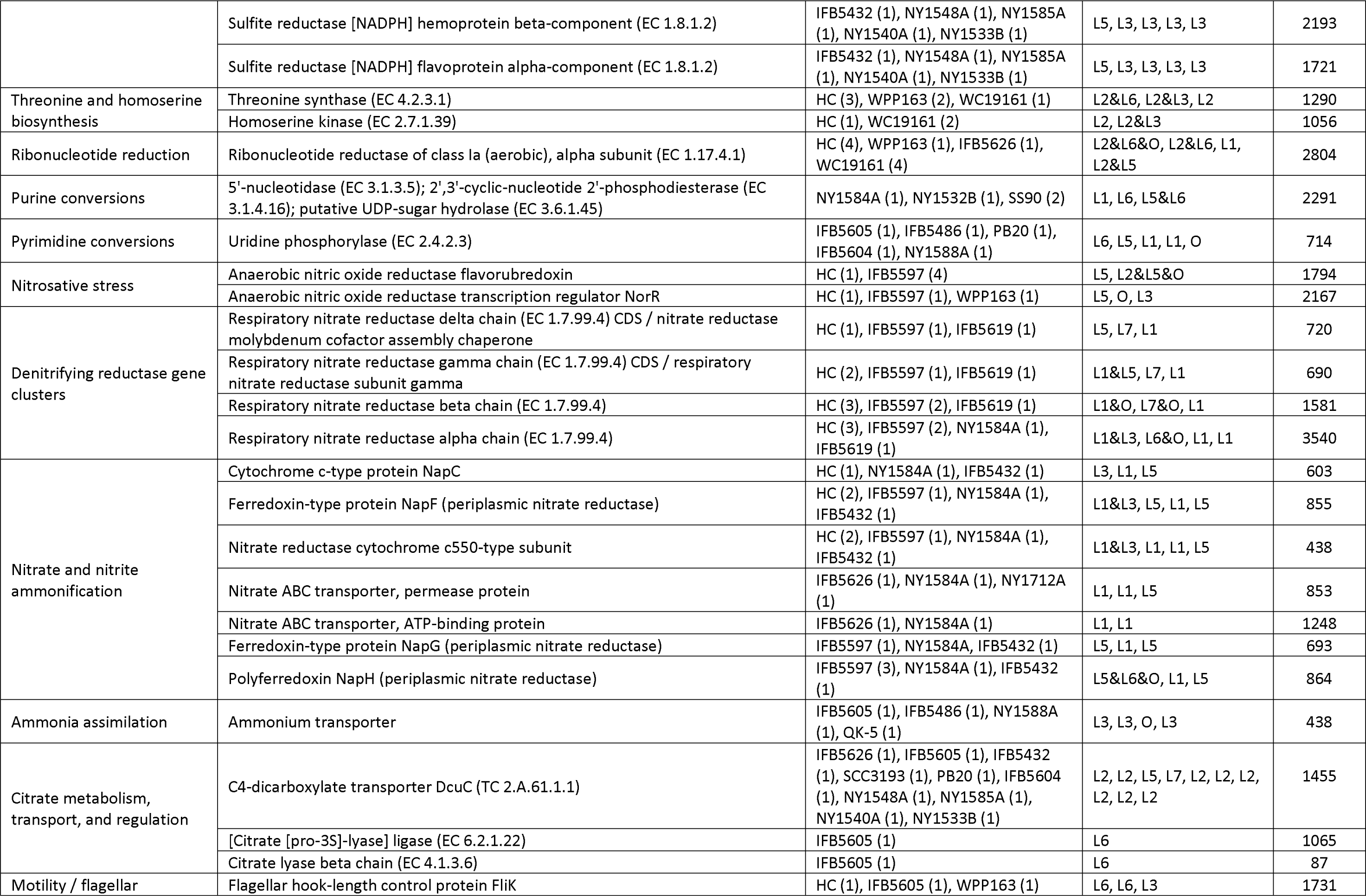
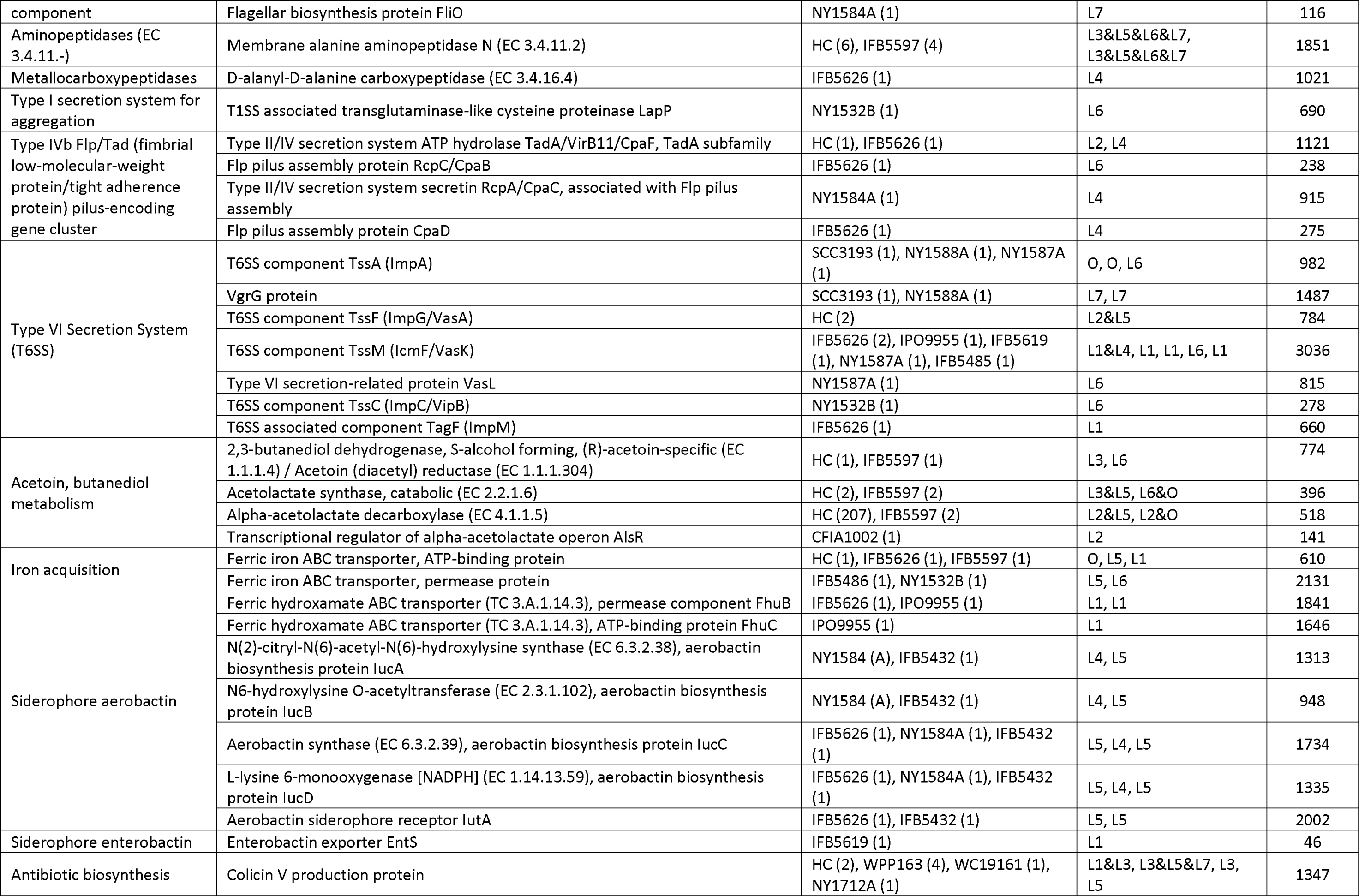
Summary of main recent recombination events predicted in the core-genome of *Pectobacterium parmentieri*. The genes where the recombination events were identified are listed per subsystem and their product name are described as annotated in the RASTtk server.

Secretion of CWDE is crucial for host colonization and disease development in soft rot bacteria (Arizala & Arif 2019; Charkowski 2018; Nykyri et al. 2012; Toth et al. 2006). In *P. parmentieri*, nine genes (*pelI*, *pehA*, *pelA*, *pelB*, *pelC*, *pelL*, *pehX*, *pelX*, *pemA*) were identified as being affected by ancestral recombination events in lineages 1, 2, 5 and 6 (**table 2**). Recombination in genes *pelA*, *pelB* and *pelC,* which encode for pectate lyases, were observed in lineages 2 and 5, while the other genes experienced recombination event in at least one lineage. Previous studies on the activity of CWDE and potato maceration efficacy have revealed significant statistical differences among *P. parmentieri* strains (Zoledowska et al. 2018a; Zoledowska et al. 2018b; Moleleki et al. 2013; Ngadze et al. 2012; Dees et al. 2017). The evidence of ancestral recombination found in our study in some of the CWDE genes might be related to the reported variation in the ability to macerate potato tissue, as well as the high differences displayed in CWDE activities within the *P. parmentieri* intraspecies.

Flagellum is another detrimental pathogenicity associated factor specially in soft rot bacteria (Charkowski et al. 2012). Mutations in flagella genes have led to a reduced virulence in *Dickeya* (Jahn et al. 2008). In *P. carotovorum* Pcc21, mutants with defective flagella biosynthesis in the genes *flgA* and *fliA* produced less than 50% of biofilm in comparison with the wild type (Lee et al. 2013). Our analysis found ancestral recombination occurring in 12 components (FliK, FlgA, FlgB, FlgC, FlgD, FlgE, FlgF, FlgG, FlgI, FliD, FliE, and FliF) of the flagellum biogenesis cluster in the lineage 1, whereas lineages 6, 2 and 3 experienced recombination in less number of genes (6 and 3 genes, respectively) (**table 2**). In addition, recent recombination events (**table 3**) were also observed within the coding sequences FliK (in strains HC, IFB5605, and WPP163) and FliO (NY1584A). These results indicate that the recombination seen in the flagellum biosynthesis cluster may be connected to the pathogen’s ability to adapt to various host conditions, given that the flagellum is a crucial component for ensuring a successful host colonization. In synchronization with the data obtained from the ClonalFrameML analysis, the ancestral and recent recombinations derived from the fastGEAR algorithm also put in evidence that other distinct and highly important pathogenicity determinants, such as genes encoding for components of the T1SS (type I secretion system), T6SS, type I secretion system for aggregation, type IV pilus biogenesis, the type IVb Flp/Tad cluster, butanediol metabolism, citrate metabolism, siderophore production, and polysaccharide biosynthesis, have undergone to homologous recombination (**tables 2-3**). These results seem to indicate that homologous recombination has a pivotal implication in the pathogenic lifestyle of *P. parmentieri*.

Notably, gene regions encoding for antimicrobial products, namely phenazine and colicin, were also affected by ancestral and recent recombination events, respectively (**tables 2-3**). Phenazines are secondary metabolites produced by bacteria and are associated with multiple roles, namely, biofilm formation, enhanced virulence, host colonization, and as weapon against other competitors like fungi or bacteria (Pierson & Pierson 2010). Colicins, on the other hand, are proteins produced by bacteria to kill other closely related members, giving them an advantage over other bacteria in the race for nutrients (Arizala & Arif 2019; Grinter et al. 2012). *P. carotovorum* Pcc21, for instance, has been reported to secrete a colicin-like bacteriocin product, Carocin D, which inhibited the growth of other *Pectobacterium* members (Roh et al. 2010). The fact that we found recombination within genes involved with the synthesis of these two antimicrobial compounds highlights the contribution of homologous recombination in the ecological fitness of *P. parmentieri*.

Moreover, our fastGEAR output showed recombination within two CRISPR-Cas adaptative immune systems of *P. parmentieri*. In the CRISPR-Cas subtype I-F system, ancestral recombination patterns were showed within the genes *cse4* in lineage 1 as well as *csy2*, and *csy3* in lineage 2 (**table 2**). In the CRISPR-Cas subtype I-E system, recent recombination events were observed within the *cas3* gene of the strains NY1584A and NY1712A (**table 3**). These findings suggest that homologous recombination seems to have participated in the evolution of the CRISPR-Cas system of *P. parmentieri*. This is consistent with earlier research that pointed recombination events spanning the genetic exchange of several *cas* gene regions (Touchon & Rocha 2010; Takeuchi et al. 2012).

### Conclusions

In the present study, we assessed the impact of homologous recombination on the core-genome of the newly, but widely spread, described species *P. parmentieri*. Clear evidence of diverse recombination patterns was found, with data indicating variations in the sizes of DNA recombinant fragments occurring randomly across the core genome of the 32 *P. parmentieri* isolates. The fastGEAR analysis identified 941 recent recombination events and 486 ancestral recombination events. The ancestral recombination data suggested that strains from lineages 1st to 6th evolved from strains of the 7th lineage. Importantly, several ancient recombinant regions (3,094) were inferred in the reconstructed ClonalFrameML genealogy tree in the main nodes, giving origin to different clades. Genes affected by homologous recombination encoded vital cell functions such as replication, transcription, metabolism, and homeostasis. Strikingly, recombination was also observed within gene loci categorized as pathogenicity determinants, including type secretion systems, iron acquisition, flagellum biosynthesis, CWDE, citrate and butanediol metabolism, type IV pilus biogenesis, etc. Recombinant fragments were also present in sequences encoding antimicrobial metabolites (phenazine and colicin) and genes of the CRISPR-Cas systems. These findings highlight the significant role of homologous recombination in shaping *P. parmentieri* genomes, influencing evolution, strain diversity, lifestyle, pathogenicity, adaptative immunity and ecological fitness of this phytopathogenic bacterium.

## Materials and Methods

### Isolate collection

In total, thirty-two *P. parmentieri* strains from different geographic origins and isolated in distinct years were used in this study (**supplementary table S1**). The seventeen complete genomes and fifteen draft genomes were retrieved from the NCBI database in FASTA format. The accession numbers used to retrieve the genomic data of the thirty-two isolates are provided in **supplementary table S1**.

### Core-genome analysis

All thirty-two genomes in FASTA format were annotated using the rapid prokaryotic genome annotation tool Prokka (Seemann 2014). The annotated genomes in GFF3 format were used as input data to carry out the pan-genome analysis in the Roary pipeline (Page et al. 2015) with a minimum BLASTP percentage identity of 90%. A multi-FASTA alignment of the previously identified core genes was generated using PRANK (Löytynoja 2014). Later, the gaps were removed from the concatenated core genome alignments using trimAl v1.3 (Capella-Gutiérrez et al. 2009) with the “-*nogaps*” parameter.

### Population genetic structure, genealogy reconstruction and inference of homologous recombination

The core-genome alignment generated with Roary for the 32 *P. parmentieri* isolates was used as input file to determine the intraspecies population structure using the hierarchical Bayesian Analysis of Population Structure (hierBAPS; Cheng et al. 2013) implemented in R (RhierBAPS; Tonkin-Hill et al. 2018). Each strain was assigned to its specific lineage genetic cluster (lineage).

Later, the generated core served to compute the maximum likelihood (ML) phylogenetic tree. The inferred phylogeny was constructed using the General Time-Reversible GAMMA (GTR-GAMMA) distribution model with 1,000 bootstraps replicates. The analysis was carried out using the RAxML-NG (RAxML Next Generation) tool (Kozlov et al. 2019). The RAxML-NG maximum likelihood (ML) tree was visualized and plotted in FigTree v1.4.4 (http://tree.bio.ed.ac.uk/software/figtree/) and ordered based on increasing order of nodes.

ClonalFrameML (Didelot & Wilson 2015) analysis was deployed to reconstruct the genealogy, infer homologous recombined fragments and polymorphisms sites within the branches and nodes across the core genome of *P. parmentieri*. The CLonalFrameML phylogeny was inferred using the previous RAxML-NG maximum likelihood tree as the starting topology tree. Recombination parameters in ClonalFrameML were calculated using the hidden Markov model based on 100 simulations (emsim=100) to refine the reliability of the analysis. Moreover, the impact of introduced recombination events relative to single-point mutations (*r/m*) was calculated for each branch in ClonalFrameML.

### Inference of recent and ancestral recombination events

The origin, number and location of recent and ancestral recombination events within the *P. parmentieri* core-genome was predicted using the novel algorithm fastGEAR, which provides information about lineages across microbial alignments and identifies recombination sites between the studied isolates as well as from external origins (Mostowy et al. 2017). A pre-specified partition file containing the number of lineages obtained in RhierBAPS was used in the fastGEAR analysis instead of the BAPS3 clustering algorithm. Once the lineages were identified in the core-alignment, recent recombinations within each lineage were detected by applying a Hidden Markov Model (HMM) approach to all subsets of strains assigned to that lineage. Ancestral recombinations were analyzed between lineages and using the same HMM approach. Regions inferred as recent recombination were removed prior to the analysis of ancestral recombination. Bayes factor >1 and >10 were used to evaluate the statistical significance of putative recent and ancestral recombination events. Additionally, the order of the thirty-two *P. parmentieri* strains shown in the recent and ancestral recombination plots was specified according to the phylogenetic tree generated in RAxML-NG. To extract the coordinates of the recombined regions and determine the annotation of genes that undergone underwent recent recombination, the next three steps, modified from Potnis et al. (2019), were followed. First, the recombination sites obtained from fastGEAR were organized into specific BED files for each *P. parmentieri* strain as well as for each node identified by ClonalFrameML (**supplementary fig. S1**). Secondly, the recombined fragments were extracted from their respective genomes using the GetFastaBed tool. Lastly, the extracted sequences were submitted to a nucleotide BLAST (BLASTn) analysis against a customized database of its corresponding *P. parmentieri* annotated core-genome. The core-genomes were annotated using the Rapid Annotation using Subsystem Technology server (RASTtk; Brettin et al. 2015). The custom BLAST analysis was performed in Geneious Prime. The same approach was followed to determine the genes affected by ancestral recombination in each of the lineages as well as to locate the recombinant fragments inferred by ClonalFrameML.

## Acknowledgements

This work was supported by the USDA-ARS Agreement No. 58-2040-9-011, Systems Approaches to Improve Production and Quality of Specialty Crops Grown in the U.S. Pacific Basin; sub-project: Genome Informed Next Generation Detection Protocols for Pests and Pathogens of Specialty Crops in Hawaii. This work was also supported by USDA NIFA Project No. HAW09706-G (Accession No. 1030040). The bioinformatics analyses were supported by NIGMS of the National Institutes of Health under award number P20GM125508.

