## Supplementary material for "Impact of homologous recombination on core genome evolution and host adaptation of *Pectobacterium parmentieri*": Table S1

**Supplementary table 1**

Detail information of the thirty-two *Pectobacterium parmentieri* genomes fetched from the NCBI GenBank database and used to assess the homologous recombination events

| **Strain name** | **NCBI accession number** | **Sequencing platform** | **Assembly method** | **Assembly level** | **Isolation source** | **Isolation year** | **Location** |
| --- | --- | --- | --- | --- | --- | --- | --- |
| RNS 08-42-1Aᵀ | NZ_CP015749 | PacBio; Illumina HiSeq | RS_HGAP v. 2.0 | Complete | Potato | 2008 | France |
| HC | NZ_CP046376 | PacBio | Other SMARTdenovo v. 1.0.0 | Complete | Potato stem | 2016 | South Korea |
| SCC3193 | NC_017845 | Roche 454 GS20; SoLiD 2; Fosmid end sequencing | Newbler | Complete | Potato stem | 1980 | Finland |
| IFB5427 | NZ_CP027260 | Illumina MiSeq | SPAdes v. JUNE-2016 | Complete | Potato associated weed | 2013 | Poland |
| IFB5441 | NZ_CP026980 | Illumina MiSeq | SPAdes v. JUNE-2016 | Complete | Potato tuber | 2013 | Poland |
| QK-5 | NZ_CP062254 | Illumina HiSeq | ABySS v. 2.0.2 | Complete | Potato | 2019 | China: QingKou |
| IFB5486 | NZ_CP026982 | Illumina MiSeq | SPAdes v. JUNE-2016 | Complete | Potato tuber | 2012 | Belgium |
| IFB5408 | NZ_CP026977 | Illumina MiSeq | SPAdes v. JUNE-2016 | Complete | Potato stem | 2013 | Poland |
| IFB5432 | NZ_CP026979 | Illumina MiSeq | SPAdes v. JUNE-2016 | Complete | Potato | 2013 | Poland |
| WC19161 | NZ_CP065946 | PacBio; Illumina NovaSeq | SMRT Link v. 5.0.1 | Complete | Potato | 2019 | China: Hohhot Inner Mongolia |
| IFB5619 | NZ_CP026985 | Illumina HiSeq | SPAdes v. MARCH-2017 | Complete | Potato stem | 2014 | Poland |
| IFB5485 | NZ_CP026981 | Illumina MiSeq | SPAdes v. JUNE-2016 | Complete | Potato | 2012 | Belgium |
| IFB5604 | NZ_CP026983 | Illumina HiSeq | SPAdes v. MARCH-2017 | Complete | Potato stem | 2014 | Poland |
| PB20 | NZ_CP065036 | Illumina MiSeq; Oxford Nanopore MinION | SPAdes v. 3.6.1 | Complete | Sewage water at the washing station of potato storage facility | 2014 | Russia: Moscow region |
| IFB5623 | NZ_CP026986 | Illumina HiSeq | SPAdes v. MARCH-2017 | Chromosome | Potato stem | 2014 | Poland |
| IFB5605 | NZ_CP026984 | Illumina HiSeq | SPAdes v. MARCH-2017 | Chromosome | Potato stem | 2014 | Poland |
| IFB5626 | NZ_PSZG00000000 | Illumina HiSeq | SPAdes v. MARCH-2017 | Scaffold | Potato tuber | 2014 | Poland |
| IFB5597 | NZ_PSZH00000000 | Illumina HiSeq | SPAdes v. MARCH-2017 | Contig | Potato stem | 2014 | Poland |
| IPO:1955 | NZ_JACGFO000000000 | Illumina HiSe | CLC NGS Cell v. 12 | Scaffold | Unknown | 2002 | Unknown |
| SS90 | NZ_QESW00000000 | Illumina MiSeq | CLC Genomics v. 10.1.1 | Contig | Potato stem | 2017 | Pakistan: Punjab |
| NY1584A | NZ_WABP00000000 | Illumina NextSeq | SPAdes v. 1.12 | Contig | Potato | 2016 | USA: New York |
| NY1532B | NZ_WABL00000000 | Illumina NextSeq | SPAdes v. 1.12 | Contig | Potato | 2016 | USA: New York |
| NY1722A | NZ_WABU00000000 | Illumina NextSeq | SPAdes v. 1.12 | Contig | Potato | 2017 | USA: New York |
| NY1712A | NZ_WABT00000000 | Illumina NextSeq | SPAdes v. 1.12 | Contig | Potato | 2017 | USA: New York |
| NY1587A | NZ_WABR00000000 | Illumina NextSeq | SPAdes v. 1.12 | Contig | Potato | 2016 | USA: New York |
| NY1585A | NZ_WABQ00000000 | Illumina NextSeq | SPAdes v. 1.12 | Contig | Potato | 2016 | USA: New York |
| NY1540A | NZ_WABN00000000 | Illumina NextSeq | SPAdes v. 1.12 | Contig | Potato | 2016 | USA: New York |
| NY1533B | NZ_WABM00000000 | Illumina NextSeq | SPAdes v. 1.12 | Contig | Potato | 2016 | USA: New York |
| NY1588A | NZ_WABS00000000 | Illumina NextSeq | SPAdes v. 1.12 | Contig | Potato | 2016 | USA: New York |
| NY1548A | NZ_WABO00000000 | Illumina NextSeq | SPAdes v. 1.12 | Contig | Potato | 2016 | USA: New York |
| CFIA1002 | NZ_JENG00000000 | Illumina HiSeq | ABySS v. MAY-2013 | Scaffold | Potato | 2007 | Canada: Alberta |
| WPP163 | NC_013421 | Roche 454 GS FLX |  | Complete | Potato | 2004 | USA: Wisconsin |

ᵀ Type strain
